# MBG: Minimizer-based Sparse de Bruijn Graph Construction

**DOI:** 10.1101/2020.09.18.303156

**Authors:** Mikko Rautiainen, Tobias Marschall

## Abstract

**Motivation:** De Bruijn graphs can be constructed from short reads efficiently and have been used for many purposes. Traditionally long read sequencing technologies have had too high error rates for de Bruijn graph-based methods. Recently, HiFi reads have provided a combination of long read length and low error rate, which enables de Bruijn graphs to be used with HiFi reads.

**Results:** We have implemented MBG, a tool for building sparse de Bruijn graphs from HiFi reads. MBG outperforms existing tools for building dense de Bruijn graphs, and can build a graph of 50x coverage whole human genome HiFi reads in four hours on a single core. MBG also assembles the bacterial *E. coli* genome into a single contig in 8 seconds.

**Availability:** Package manager: https://anaconda.org/bioconda/mbg and source code: https://github.com/maickrau/MBG

## 1 Introduction

De Bruijn graphs have been used for a long time in sequence analysis for purposes such as genome assembly [10, 1, 16, 4] and error correction [13, 7, 11]. Sparse de Bruijn graphs [17] are a form of de Bruijn graph which use only a subset of k-mers and so reduce runtime and memory use. Minimizer winnowing [14, 12] is a method of selecting a subset of k-mers from a sequence. Minimizer winnowing has been applied to building sparse de Bruijn graphs [3]. Recently, HiFi reads [15] have reached read lengths of thousands of base pairs with error rates comparable or superior to shotgun sequenced short reads. The combination of long read lengths and low error rates makes de Bruijn graphs an attractive idea for HiFi reads and might enable hybrid methods for genome assembly and error correction to use HiFi reads. However, current tools do not scale for building de Bruijn graphs with k-mer sizes in thousands.

### Contributions

We have implemented the tool MBG (**M**inimizer-based sparse de **B**ruijn **G**raph) for constructing sparse de Bruijn graphs. MBG selects k-mers by minimizer winnowing [14] and builds the graph from those k-mers. This approach has previously been used in the ntJoin scaffolder [3] for building graphs from assembled contigs to scaffold assemblies.

MBG can construct graphs with arbitrarily high k-mer sizes, and we show in the experiments that k-mer sizes of thousands of base pairs are practical with real HiFi read data. MBG outperforms existing de Bruijn graph construction tools in runtime, with a runtime of only a few hours on a single core for constructing a graph of 50x coverage whole human genome HiFi reads.

## 2 Methods

We give a brief overview of the implementation here with detailed explanations of the individual steps in Appendix A. Since most errors in HiFi reads are homopolymer run length errors [15], the input reads are homopolymer compressed by collapsing homopolymer runs into one character. A rolling hash function [8] is then used to assign a hash value to each k-mer. Minimizer winnowing [14] is then used to select the smallest k-mer in each window. The selected k-mers are compressed by hashing them into 128-bit integers, which form the nodes of the minimizer graph. Edges are added whenever two minimizers are adjacent to each other in the reads. Transitive edges caused by sequencing errors are cleaned. Non-branching paths of the graph are then condensed into unitigs. Finally, the 128-bit hashes are replaced with their base pair sequences, and homopolymer runs are expanded. The graph is then written in the GFA format [6].

## 3 Results

We built sparse de Bruijn graphs using HiFi read data. Details of the experimental setup are in Appendix B. Table 1 shows the results.

**Table 1:** Experimental results

| Dataset | Tool | k | w | CPU-time | Memory (Gb) | N50 |
| --- | --- | --- | --- | --- | --- | --- |
| *E. coli* | BCalm2 | 61 | - | 0:01:49 | 1.1 | 1025 |
|  |  | 91 | - | 0:04:10 | 1.6 | 1080 |
|  |  | 127 | - | 0:07:20 | 2.0 | 1212 |
| *E. coli* | MBG | 45 | 1 | 0:01:46 | 2.8 | 60479 |
|  |  | 41 | 10 | 0:00:32 | 0.5 | 59657 |
|  |  | 35 | 20 | 0:00:20 | 0.3 | 57134 |
|  |  | 31 | 30 | 0:00:18 | 0.2 | 34101 |
|  |  | 69 | 1 | 0:01:44 | 3.1 | 82820 |
|  |  | 65 | 10 | 0:00:34 | 0.6 | 82810 |
|  |  | 59 | 20 | 0:00:23 | 0.4 | 73682 |
|  |  | 55 | 30 | 0:00:20 | 0.3 | 78679 |
|  |  | 95 | 1 | 0:02:13 | 3.4 | 117784 |
|  |  | 91 | 10 | 0:00:39 | 0.7 | 117782 |
|  |  | 85 | 20 | 0:00:25 | 0.4 | 125638 |
|  |  | 81 | 30 | 0:00:19 | 0.3 | 117643 |
| *E. coli* | MBG | 501 | 500 | 0:00:10 | 0.12 | 177653 |
|  |  | 1001 | 1000 | 0:00:09 | 0.13 | 698111 |
|  |  | 1501 | 1500 | 0:00:09 | 0.14 | 1517634 |
|  |  | 2001 | 2000 | 0:00:08 | 0.14 | 4639237 |
|  |  | 2501 | 2500 | 0:00:08 | 0.14 | 4644046 |
|  |  | 3001 | 3000 | 0:00:08 | 0.14 | 4090727 |
|  |  | 3501 | 3500 | 0:00:07 | 0.14 | 392371 |
| HG002 | MBG | 501 | 500 | 4:10:28 | 123.7 | 2012 |
|  |  | 1001 | 1000 | 4:38:40 | 132.6 | 4501 |
|  |  | 2001 | 2000 | 4:07:22 | 138.5 | 12095 |
|  |  | 3001 | 3000 | 3:57:59 | 137.3 | 23104 |
|  |  | 4001 | 4000 | 3:35:03 | 120.4 | 26736 |
|  |  | 5001 | 5000 | 2:06:52 | 68.9 | 20699 |
|  |  | 501 | 100 | 6:50:21 | 269.0 | 1784 |
|  |  | 1001 | 200 | 6:16:38 | 244.2 | 3749 |
|  |  | 2001 | 400 | 6:01:18 | 250.0 | 8669 |
|  |  | 3001 | 600 | 6:05:17 | 243.9 | 15085 |
|  |  | 4001 | 800 | 3:55:50 | 245.3 | 23868 |
|  |  | 5001 | 1000 | 4:47:44 | 244.1 | 33649 |

### Comparison to existing tools

We compared MBG to BCalm2 [2] for building graphs using HiFi reads of *E. coli*. Note that N50 is not directly comparable between MBG and BCalm2 since the homopolymer compression step removes most errors and therefore greatly improves N50. BCalm2 uses less memory than MBG with *w* = 1, but for *w* = 10 and higher MBG uses less memory. MBG is faster for all tested values of *k* and *w* except *k* = 91 and *w* = 1 which is slower than BCalm2 with *k* = 61. With *w* = 30 MBG is an order of magnitude faster than BCalm2. With *k* = 2501 and *w* = 2500, MBG assembles *E. coli* correctly into a single contig in 8 seconds on a single core.

### Whole human genome HiFi

We ran MBG on whole human genome HiFi data from the individual HG002. Runtime is between 2 and 7 hours on a single core with all parameter sets, showing that MBG is fast and scales to large values of *k*. The limitation on increasing k even higher is the error rate and read length of the HiFi reads.

## 4 Conclusion

We have implemented MBG, a tool for building sparse de Bruijn graphs from HiFi reads using minimizer winnowing. The sparsification enables MBG to run orders of magnitude faster than tools for building dense de Bruijn graphs. MBG uses a novel method to compress long k-mers to constant sized hashes and enables k to scale arbitrarily high.

MBG can quickly build de Bruijn graphs of mammalian sized genomes, with runtimes ranging from 2 to 6 hours on a single core. The memory use currently prevents MBG from being ran on mammalian datasets on laptops and desktop computers. However, MBG fits comfortably in the RAM of most computing servers. MBG enables small genomes such as *E. coli* to be assembled in a few seconds and mammalian genomes in a few hours.

## A Methods

### Homopolymer compression

Most errors in HiFi reads are homopolymer run length errors [15]. The sequences are first homopolymer compressed by collapsing homopolymer runs into one character, reducing the error rate by an order of magnitude. The lengths of the homopolymer runs are stored so that the original sequence can be reconstructed at the end.

### Minimizer winnowing

MBG uses the rolling hash function from ntHash [8] to assign hash values to each k-mer of the input reads. The runtime of the rolling hash function is independent of k-mer size. In practice minimizer winnowing is the performance bottleneck of MBG, so we chose the ntHash method since it is the fastest hash we are aware of.

Minimizer winnowing [14] is then applied to the k-mers given their hash values. The smallest k-mer in each window is selected for later processing. Selected k-mers which appear in the input data fewer times than a user given k-mer abundance cutoff are also discarded. Since the density of random minimizers is 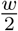 [14], a window size of *w* will on average lead to a 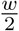-fold sparsity of selected k-mers.

### Compressing arbitrary sized k-mers by hashing

The selected k-mers are compressed by hashing them into 128-bit integers. The 128-bit hashes are then used as the nodes of the graph. We used the c++ standard library’s string hash function for building the hash. For a string *s*, the lower 64 bits of the hash are taken from the standard library hash of 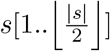 and the upper 64 bits from the hash of 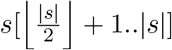. Note that hash quality is very important in this step. Since a hash collision would lead to two different sequences being represented by the same node, every k-mer must result in a unique hash to ensure correctness of the resulting graph.

### Hash collisions

Given 128-bit random hashes, it is reasonable to assume that there are no hash collisions. To estimate the probability of a hash collision, the birthday paradox can be used. Given *n* k-mers to hash, and the size of the hash space *d* = 2^128^, the probability of collision can be approximated with 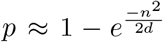. As of 1st July 2020, the size of the SRA database is 42441459655506377 base pairs. If the entire database were concatenated to one string and all of its k-mers for one k were hashed, there would be less than 4.3 * 10^16^ k-mers to hash. Applying the approximation of the birthday paradox to this number of k-mers gives a hash collision probability of 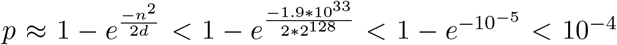 for hashing the entire SRA database. The probability of hash collision for any realistic dataset is therefore negligible assuming a random hash function. In addition, MBG checks for hash collisions during runtime. We have not seen a hash collision so far.

### Transitive edge cleaning

Because the minimizers are sampled from a window, a sequencing error outside of a k-mer can affect whether the k-mer was chosen. That is, an error within a window but outside of the chosen k-mer in the error-free window can cause a different k-mer to be chosen in the error-containing window.

Figure 1 illustrates the problem. The error-free sequence has chosen k-mers *x*_1_, *x*_2_, and *x*_3_. The second sequence has a sequencing error outside of *x*_2_ but covered by each window that selected *x*_2_, causing *x*_2_ to not be selected, and instead minimizers *x*_4_ and *x*_5_ are selected. If the minimizers were used as-is, the graph would have an extra edge *x*_4_ → *x*_4_, and the correct edge *x*_2_ → *x*_4_ would be missing.

**Figure 1:**
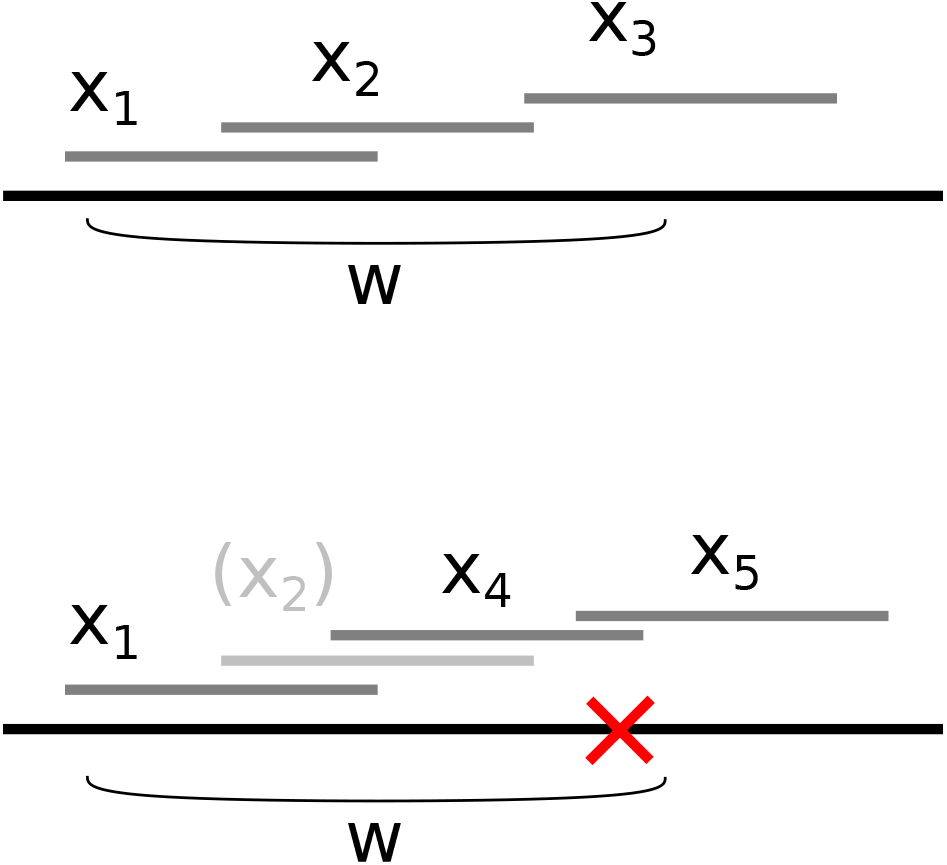
An illustration of the transitive edge problem. The top sequence (solid black line) has no errors and three k-mers, *x*_1_, *x*_2_ and *x*_3_, are selected from it. The area marked by *w* is one window, from which *x*_2_ was selected in the error-free sequence. The bottom sequence (solid black line) has a sequencing error (red cross). Due to the sequencing error, k-mer *x*_4_ is selected from window *w* instead of *x*_2_. The k-mers selected from the bottom sequence are *x*_1_, *x*_4_ and *x*_5_. Even though the bottom sequence contains *x*_2_ without errors, *x*_2_ is not selected.

To solve this, we look at all edges connecting minimizers. We build an *edge sequence* of the two adjacent minimizers. Given k-mers *m*_1_ with sequence *σ*_1_ and *m*_2_ with sequence *σ*_2_, and an overlap of *b* base pairs between them, the edge sequence is defined as *s* = *σ*_1_ + *σ*_2_[*b..k*], that is, the concatenation of the two k-mers, taking into account not to duplicate the shared sequence in the overlap. Then, we check all k-mers in the edge sequence. If a k-mer *m*_3_ inside the edge sequence was selected as a minimizer during minimizer winnowing, we mark the edge (*m*_1_, *m*_2_) as transitive, and add the edges (*m*_1_, *m*_3_) and (*m*_3_, *m*_2_) if they were not already present. Finally, we remove all transitive edges and transfer their read coverage to the replacement edges. In the example in Figure 1, this would remove the edge *x*_1_ → *x*_4_, add the new edge *x*_2_ → *x*_4_, and add the coverage of the removed edge *x*_1_ → *x*_4_ to the edges *x*_1_ → *x*_2_ and *x*_2_ → *x*_4_.

Checking if a k-mer was selected as a minimizer takes *O*(*k*) time and checking all k-mers in all edges would then take *O*(*mk*^2^) time for *m* minimizers. We improve the speed in practice by first using the rolling hash from minimizer winnowing to limit which k-mers to check. The hash values of the selected k-mers are stored, and then a k-mer within an edge sequence is checked only if its hash value exists in the stored hash values. This check can be done in *O*(*mk*) time for all k-mers in all edges. Empirically, more than 99.99% of k-mers that pass the rolling hash check are also selected k-mers. This does not affect the theoretical runtime of the algorithm but in practice it leads to a significant speedup.

### Graph construction

The 128-bit hashes are used as the nodes of the graph. Edges are added whenever two hashes are adjacent to each other in a read. The constructed graph is then processed by condensing non-branching paths into *unitigs*. After this, unitigs are filtered based on a user given unitig abundance cutoff. Unitigs whose average coverage is less than the cutoff are discarded. In addition, edges whose coverage is less than the cutoff are also discarded. After unitig and edge removal the non-branching paths are again condensed into unitigs. The 128-bit hashes are transformed back to base pair sequences and homopolymer runs are decompressed. Finally, the graph is written in GFA format [6].

### Storing sequences

The base pair sequences of the selected k-mers are stored in memory as a store that contains a list of contiguous blocks. When a k-mer is added to the store, if the k-mer has overlap with the most recently added k-mer, the non-overlapping part is appended to the contiguous block. That is, the overlapping part is only stored once. If the k-mer does not overlap with the most recently added k-mer, a new block is started. The lengths of the homopolymer runs are stored similarly using 16-bit integers. In practice this means that adjacent k-mers from the same read can be stored efficiently without duplicating the overlapping sequences, and moving from one read to another will almost certainly start a new contiguous block.

### Homopolymer run length consensus

When storing the sequences, homopolymer run lengths are also stored. Each stored base pair also has a sum of run lengths *s* and a base pair count *c*. When a k-mer is read from the input, for every base pair in the k-mer, the base pair count *c* of the associated base pair is incremented by one, and the homopolymer run length of the k-mer is added to the sum of run lengths *s*. The run length consensus of each base pair is taken from the average 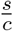 rounded to the nearest integer. The run length consensus can optionally be disabled to reduce memory use, which is intended for the case when the input reads are already homopolymer compressed.

### Runtime

Assuming no sequencing errors and given a genome size g, genomic coverage *c*, k-mer size *k* and window size *w*, the number of selected minimizers is *m* with 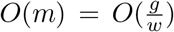 assuming the minimizer winnowing hash is random. The runtime of minimizer winnowing is *O*(*gc*). Hashing the selected k-mers is 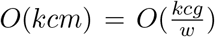. Cleaning transitive edges requires 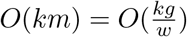 for the selected minimizers, and 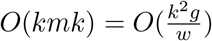 for k-mers which share their rolling hash value with a selected minimizer. Graph construction is 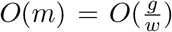. In total the runtime is 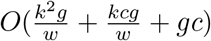. In practice the 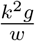 term has a tiny constant factor and the runtime is dominated by the 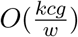 term.

Assuming no sequencing errors and a constant read length *r* > *k* + *w*, the memory use of MBG is 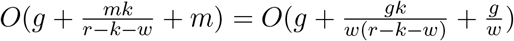. In practice increasing *w* reduces memory use significantly.

## B Experimental setup

We used MBG version 1.0.1 from Bioconda. We used BCalm2 version 2.2.3 from Bioconda. All experiments were ran on a computing server with 48 Intel(R) Xeon(R) E7-8857 v2 CPUs and 1.5Tb of RAM. BCalm2 was given one thread in the command line invocation, and MBG is single threaded. Runtime and memory use was measured with “/usr/bin/time -v” in all experiments.

### Comparison to existing tools

We compared MBG to BCalm2 [2] for building graphs. We used HiFi data from *E. coli*^1^, containing 290x coverage HiFi reads. We randomly downsampled the reads to 29x coverage. We varied the window size parameter *w* for MBG from 1, resulting in a de Bruijn graph, to 30, sparsifying the k-mer set by a factor of 15 on average. The k-mer abundance threshold was set to 3 for BCalm2, and the unitig average abundance threshold was set to 3 for MBG.

Due to the average density of random minimizers of *w*/2, and the homopolymer compression reducing the average length of sequence by 1/4, the results for a de Bruijn graph with k-mer size *k_DBG_* are most closely comparable to a sparse de Bruijn graph with 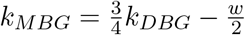. We tried different values of *k_MBG_* and w which result in similar graph quality as predicted by the above equation, and which match the *k_DBG_* given to BCalm2.

We limited *k_DBG_* in the comparison to at most 127 since that was the highest *k* supported by the version of BCalm2 we used. We also included one block with parameters more suitable for MBG.

Since the N50 of the *k* = 2001 and *k* = 2501 graphs matches the *E. coli* genome size, we evaluated their correctness by running QUAST [5] on the *E. coli* K-12 substring MG1665 reference genome^2^, and a de novo HiCanu [9] assembly of the same HiFi reads. The results were the same for *k* = 2001 and *k* = 2501 graphs produced by MBG. When compared to the reference genome, QUAST reported 8 misassemblies for both the MBG contigs and the HiCanu de novo assembly, all at the same locations. On the other hand the MBG contigs and the HiCanu de novo assembly were structurally consistent with each others. We suspect that the difference is due to the sequenced strain having differences to the strain used for constructing the reference genome. The MBG contigs had a substitution error rate of 7.8 * 10^-6^ for *k* = 2501 and 6.7 * 10^-6^ for *k* = 2001, and an indel error rate of 5.0 * 10^-4^ for *k* = 2501 and 4.5 * 10^-4^ for *k* = 2001. Nearly all of the errors are incorrect homopolymer run lengths. With homopolymer compressed reference and contigs, the error rate drops to just 3 substitution and 3 indel errors over the entire *E. coli* genome, for a total error rate of 1.8 * 10^-6^.

### Whole human genome HiFi

We ran MBG on whole human genome HiFi data from the individual HG002. We used HiFi reads from the Human Pangenome Reference Consortium HG002 data freeze v1.0 [15]^3^. The reads contain 50x coverage HiFi reads with sizes ranging from 15kbp to 25kbp.

## Footnotes

1 SRA accession number SRR10971019

2 GenBank accession U00096.2

3 Libraries m64012_190920_173625, m64012_190921_234837, m64015_190920_185703, m64015_190922_010918, m64011_190712_225711, m64011_190726_220327

## Notes

### Competing Interest Statement

The authors have declared no competing interest.

